# An individual recognition system for Hartmann’s mountain zebras (*Equus zebra hartmannae*)

**DOI:** 10.1101/2025.03.07.641993

**Authors:** L. Morris Gosling

## Abstract

Individual-based techniques are increasingly used for conservation management because of their predictive value for small populations. Mountain zebras are suitable for this approach because their stripe patterns are highly variable and individually distinct. A novel system based on sorting manually coded stripe variants at defined positions on the body has been developed for individual recognition in Hartmann’s mountain zebra (*Equus zebra hartmannae*) and is described here as an alternative to computerised pattern recognition approaches. Camera trapping at water holes and/or normal digital photography allows representative sampling of wild populations of this water-dependent equid. This approach can be used in areas which are partly inaccessible for ground survey and avoids the cost and disturbance of physical capture for artificial marking. With a comprehensive network of camera traps, virtually all individuals in a population can potentially be identified at low cost. Recognition of an individual using this system took 56+/-15 seconds (mean +/-SD) in a sample of good quality photographs from an ID library of 3,156 individuals. ID libraries at 6 main study sites where mountain zebra are routinely monitored contain over 12,000 coded individuals with about 7,800 known to be alive. The data obtained are being used for a variety of individual-based approaches including simple population monitoring using retrospective census, mark-recapture estimates, life-history analysis, individual-based modelling and studies of social behaviour.

## INTRODUCTION

Individual-based approaches provide a rigorous basis for basic and applied objectives in population and behavioural ecology. For example, known individuals can be enumerated or used in mark-recapture estimates and they are useful in behavioural ecology and individual-based population models (e.g. Grimm & Railsback, 2005), not least because they reflect the level of selection in natural populations. They are particularly useful in support of the conservation of endangered species where the survival and reproduction of small numbers of animals may be vital for population viability. When populations are small or hard to sample for other reasons, higher level demographic approaches based on birth and death rates may be inaccurate or inappropriate.

The drawback of the individual-based approach is that initial recognition and the accumulation of individual records is very time consuming. The two main techniques used are to manually code distinctive aspects of an animal’s appearance or to use computer-based pattern recognition. Both approaches have been used for zebras, notably the coding system used for Grevy’s (Ginsberg, 1987), and the increasingly powerful computer-based pattern recognition systems on plains and Grevy’s zebra (Foster et al., 2006; ‘StripeSpotter’, Lahiri et al., 2011; ‘Hotspotter’, Crall et al., 2013; Stennett et al., 2022). Pattern recognition is best known for its use in human fingerprinting and advanced techniques are rapidly developing in animal studies. Other studies of zebra use direct visual comparison of animals either in the wild or in photographs with individual reference photographs (e.g. Penzhorn, 1984) and this is effective when the population is small.

The technique described here for research on mountain zebras is to manually code stripe variation at defined stripe positions and then use a spreadsheet filter function to compare a candidate individual with these characteristics in an ID library. All the material considered is based on photographs either from camera traps or normal photography. The stripe code technique is compared with other approaches, especially computerised pattern recognition and strategies are discussed to deal with such matters as the detection of false negatives (falsely concluding that an animal is not known when in fact it is already in the ID library) and rules for the recruitment of new individuals.

The techniques reported were developed as part of the Mountain Zebra Project at the Namibian Nature Foundation. Work started in 2005 on the mountain zebra population in Gondwana Canyon Park and the adjacent Ai-Ais-Richtersveld Transfrontier Park in southern Namibia and has since been extended to NamibRand Nature Reserve, BüllsPort Guest Farm, the central Namib (in fact the Naukluft extension of the Namib-Naukluft NP and nearby private properties), Etosha NP and the Hobatere Concession Area. At the time of writing, in 2025, ID libraries at 6 main study sites where mountain zebra are routinely monitored contained over 12,000 coded individuals with about 7,800 known to be alive.

Mountain zebras in Namibia are Hartmann’s mountain zebra (*Equus zebra hartmannae*). Recent genetic studies (Moodley & Harley, 2005) do not support the specific status of Hartmann’s but the possession of unique alleles confirm their genetic separation from Cape mountain zebra (*E*.*z*.*zebra*) and they can be regarded as a sub-species. Collectively, mountain zebra, *Equus zebra*, are an endangered species (IUCN Red List Category Vulnerable) and Hartmann’s mountain zebra are a ‘Specially protected species’ in Namibia; further details are given by Novellie et al. (2002), Penzhorn (2013) and Gosling et al. (2019).

Hartmann’s mountain zebra are locally abundant in Namibia and may come into conflict with livestock farming. They are frequently reintroduced to private and public land for consumptive or non-consumptive use, including trophy hunting and game viewing (Barnes et al., 2009), and efficient counting and monitoring techniques are important for conservation management.

### COLLECTING PHOTOGRAPHS OF INDIVIDUALS

All identification in the work reported here was from photographs obtained from camera traps or conventional photography in the field. It is possible to use the technique directly with live animals but photography is preferable because it allows a permanent record and potential audit of the system. Mountain zebra are water dependent and so animals can be intercepted using camera traps at water sources. A large variety of such cameras are available and various models including Reconynx, Cuddeback, Covert,BuckEye, Ranger and Browning were used; models were selected for maximum image clarity. Mountain zebras often drink at night, particularly where they are hunted or where there is other human disturbance and infra-red (IR) light sources in cameras allows night time monitoring. Zebra approach water along well-defined trails and cameras can also be placed alongside them to photograph the side of the animal as it walks past. However, good quality images (which are essential for individual recognition) are best obtained when animals are stationary and most images taken of moving animals with IR flash are motion-blurred. Thus it is best to place cameras at water holes. These positions yield more images than those next to trails and while many animals are photographed at angles that are not useful for individual recognition the large numbers of images obtained mean that adequate samples are soon obtained. While difficult to test, this approach probably meets the sampling requirements of any population estimates in that the occurrence of an identifiable individual is random with respect to the occurrence of the individual in the population.

The placement of camera traps and their protection from various potential sources of damage varies according to the information required. There are also various trade-offs: for example a camera placed a little way from the water hole (say 15-20 metres) is more likely to detect entire social groups but less likely to obtain images of sufficient quality for individual recognition. Since the latter is the priority for individual-based studies, I generally placed cameras at about 7-8 metres from the place where spoor suggested most zebra were drinking. Individual recognition is time-consuming and, for purposes of population dynamics, I usually restricted IDs to the right side of the body. Cameras were thus placed to one side of the water hole to photograph the right sides of animals drinking and waiting to drink.

The camera traps used alternate between normal photography in daylight and IR at night. Cameras (Cuddeback) using white flash at night were used initially but these probably cause more disturbance than IR flash and were subsequently avoided. In my experience, mountain zebra can detect all cameras at night either by visual components of the flash, their noise or both. Some relatively undetectable (‘dark flash’) cameras exist but their IR flash usually has a reduced range. Mountain zebra are cautious and are usually disturbed by recently installed cameras. Habituation can take weeks or even months and varies between individuals. This change in behaviour potentially biases population sampling and any investigation must take into account the period needed for habituation. For this reason cameras are best left in the same place for long periods.

Cameras are damaged by a variety of animals. Hyenas (*Crocuta crocuta*) and baboons (*Papio ursinus*) often invstigate them and variously destroy and dismember them. Oryx (*Oryx gazella*) beat them with their horns and other animals, including mountain zebra, used them as rubbing posts. A number of things can be done to reduce such damage but most important is to fix the camera firmly to a metal or wooden post , protect the base of the post with large rocks and/or tree branches (to discourage animals from standing nearby and rubbing) and use a metal protective box. Some manufacturers produce custom-made metal boxes but when these are not available, boxes can be made by local metal workers.

In addition to camera trapping mountain zebra can be photographed using conventional photography in the field. The advent of high-definition digital photography has transformed this activity for field biologists: images can be immediately checked to see if the quality is adequate for individual recognition or if further attempts are needed. I used a 500mm lens and photographs of animals at distances of less than about 3-500m were often useful. Such photographs give information about ranging behaviour that extends the information obtained from water holes and they sometimes confirm details of social group membership which are often obscure in camera trap photographs.

### SAMPLING MOUNTAIN ZEBRA POPULATIONS

Many aspects of sampling are the same as for other populations of large terrestrial vertebrates but mountain zebra pose additional problems by living in difficult broken terrain. In areas where they have been hunted they have very long flight distances from people and/or cars, sometimes more than a kilometre. They can be sampled using road-transects (preferably using distance-sampling; Buckland et al., 2001) where adequate coverage of roads and tracks and skilled observers are available. However, difficulties of access and cryptic behaviour (such as running into gullies at the sound of approaching cars) can lead to violation of the assumptions of such survey techniques. Where these problems occur they can be overcome by individual recognition and mark-recapture techniques.

All study sites have some artificial waterholes (with solar or wind powered pumps) created to provide water for wildlife, in addition to natural waterholes. It is possible to ensure that entire populations of mountain zebra are sampled where the water holes used are known and monitored by camera traps. Whether or not a population is being adequately sampled can be determined by recognising the individuals that use particular water holes and calculating the connectivity between them. Connectivity between two water holes can be expressed as the number of individuals that occur at both expressed as a percentage of the number that occur at either, during a defined time period. If two adjacent water holes prove to have no animals in common there a risk of an un-sampled sector of the population between them. If the overlap is large, say over 70%, the effort applied at one of the water holes may be better deployed to less intensively sampled areas. The aim is to establish a network of camera traps with a reasonable degree of overall coverage so that the entire population is sampled. Optimum spacing between camera traps varies between habitats and areas but calculated connectivity values in the study areas considered here suggest that spacing between cameras of 5-10 km is a useful starting point for long-term studies in the study areas discussed here.

Mountain zebras live in large ranges (Joubert,1972; Penzhorn,1982a) and are sometimes extremely mobile. Except where fences are very well maintained, populations generally move more or less freely between landholdings including protected areas and private farms. As an ID library is accumulated it soon becomes clear that the number of animals using a particular site (such as a water hole, or land holding) is greater than the number present at any one time. This larger number can be regarded as the *‘source population’* for a particular site, that is, the animals that use the area at some times of their lives or at particular seasons but are not present at any one time. The source population can be determined by enumeration (a list based on the ID library) by retrospective census (Gosling et al, 1981) and, when sampling is constant, by extrapolation from historical capture rates of known individuals. The animals present at one time can be estimated by mark-recapture or, where possible, road transect sampling. The distinction between a source population and an instantaneous estimate is important for conservation management because a population may be larger than previously believed (and thus genetically more viable) and because the threats that face a protected population will include factors that occur throughout the entire range, including those outside a protected area, such as poaching or commercial game-capture.

### THE RECOGNITION SYSTEM

It is possible to identify individual mountain zebra by accumulating a reference library of images and visually comparing new images with each in turn. Focusing on variable areas such as features of the shoulder or rump speeds up this process. It is helpful to identify a rare stripe variant in one location on the new image and to look for this specifically using a presence or absence approach as the reference library is scanned. However, this technique is only practical when the population is quite small, say, less than about 70 animals. Above this number, a faster identification system becomes increasingly important to reduce the time spent scanning images.

Stripe patterns are highly variable in mountain zebras. Potentially any part of the body can be used in an individual recognition system including the bars down the legs, the neck and face. However, the main stripes on the shoulder, body and flanks are most conspicuous and these form the basis of the scheme outlined here. A key problem in any coding system is where to start; the zero point. Two starting points are used here, the ‘gridiron stripe’ which descends obliquely forwards from the gridiron pattern over the base of the tail (Fig.1) and the ‘shoulder stripe’, a vertical stripe above the foreleg. The gridiron stripe itself, the ‘rump stripes’ below it on the flank, and the ‘body stripes’ anterior to it, are highly variable and thus potentially useful. They are also the largest stripes on the body and this is helpful since image quality is variable in both normal photography and camera traps. A second system is based on the ‘shoulder stripe’, and the ‘rib stripes’ posterior to it. Having two separate systems, based on the rump and shoulder is useful to provide additive variation, as alternatives when the image under consideration is incomplete and as a check when trying to decide if an unrecognised animal is in fact new. This check thus aims to avoid false negatives, that is, concluding that an animal is not known (i.e. new) when in fact it is already in the ID library.

**Figure 1.**
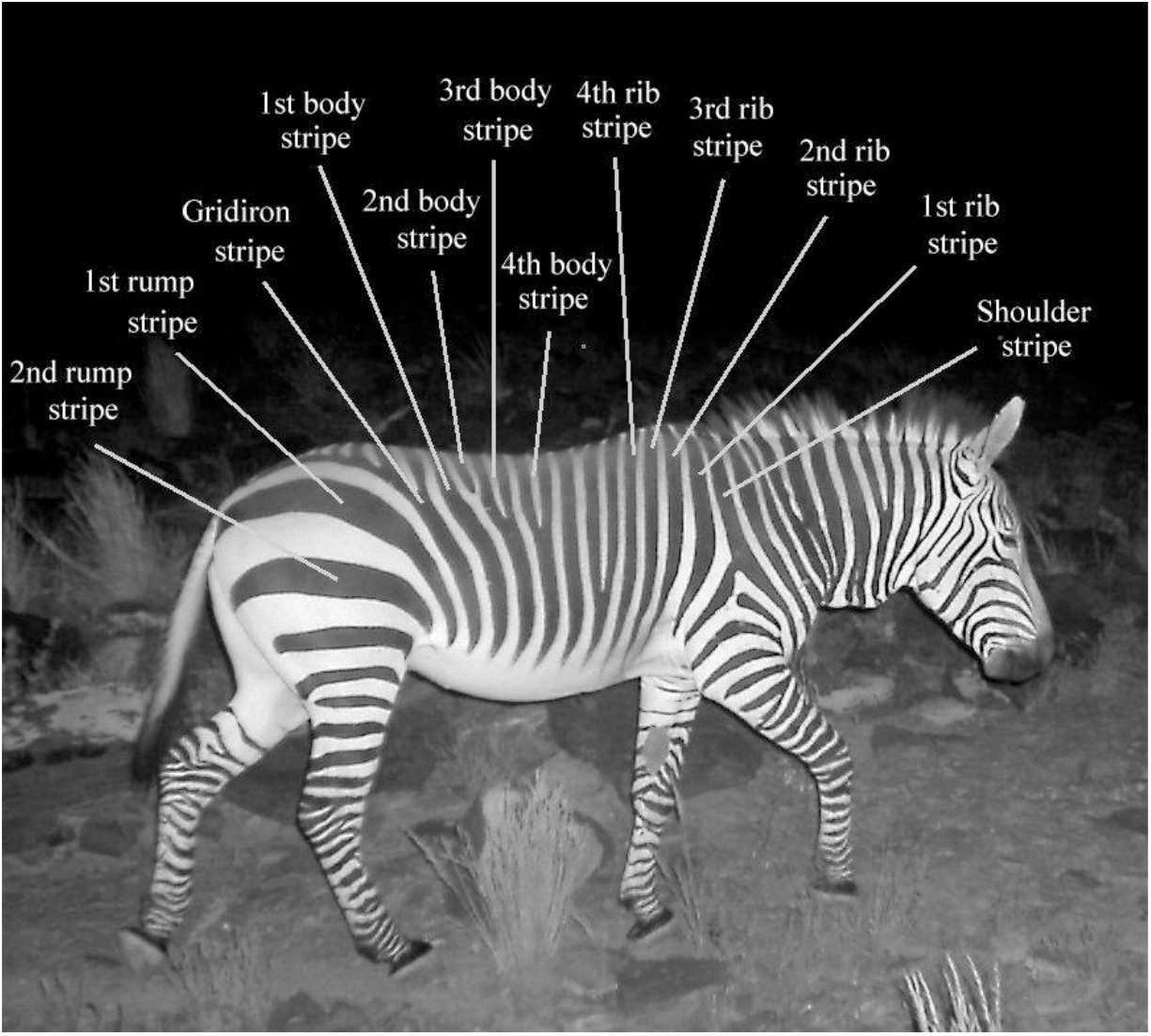
Names of stripes used in the basic individual recognition system. The gridiron stripe and shoulder stripe can be used as separate origins for coded sequences, partly to make use of incomplete images. For this individual (ZR035m from Gondwana Canyon Park), following the terminology in the text for stripe variants, the gridiron stripe is ‘simple’ and the four body stripes from left to right are: ‘simple’/’short’/ ‘Y’ shaped’/ ‘Y’ shaped; all three numbered rib stripes and both rump stripes are ‘simple’. When individuals have been provisionally identified using a key the identification can be visually confirmed using variation from another part of the body such as the details of stripes on the legs or the face. Photo © L. M. Gosling.

The individual recognition technique is based on coding stripe variants at locations on the body described above. When a new individual is found the code for each stripe is entered into an Excel database consisting of a row of characters for each individual (see Table 1). Thus body stripes can be either ‘simple’ (a simple linear shape), joined to an anterior stripe, ‘branched’ (a single dichotomous branch with the branch distal to the back of the animal), ‘Y’ shaped (with the top part of the ‘Y’ proximal to the back of the animal), ‘V’ shaped, ‘tree’ like (related to a ‘Y’ but with more than two branches), ‘spurred’ (a short side projection on a simple stripe), ‘forked’ (a small branch near the tip) or ‘short’ (distinctly shorter than neighbours at the distal end). ‘Islands’ are parts of stripes that are detached from their origin on the back of the animal and their variety is defined by their position relative to other stripes (for example beneath the gridiron stripe or beneath the first rump stripe). There is a lot of variation within each of these categories; ‘trees’ for example have variable numbers of branches and points of origin. But defining too many variants would make the operation of the system impractical and tests show that the number of variants distinguished here is sufficient. Complete lists of stripe variants and codes are given in Appendix 1.

**Table 1.** The arrangement of the Excel spreadsheet employed to record codes for the 13 characters used to identify individual mountain zebra. The example shown is the individual shown in Fig. 1 ,ZR035m (that is, zebra/right side/number 35/male) from Gondwana Canyon Park. ‘G’, ‘S’, ‘Is’ and ‘Sh’ are used as prefixed for ‘gridiron’, ‘stripe’, ‘island’ and ‘shoulder’ respectively.

| ID number | Grid stripe | 1st Rump | 2nd rump | Islands | 1st body | 2nd body | 3rd body | 4th body | Shoulder | 1st rib | 2nd rib | 3rd rib | 4th rib |
| --- | --- | --- | --- | --- | --- | --- | --- | --- | --- | --- | --- | --- | --- |
| ZR035m | G13 | S13 | S13 | Is1 | S13 | S04 | S08 | S08 | Sh3 | S13 | S13 | S13 | S13 |

Some variants of the gridiron stripe are shown in Fig. 2 and a complete list of variants, together with their codes is in Appendix 1.

**Figure 2.**
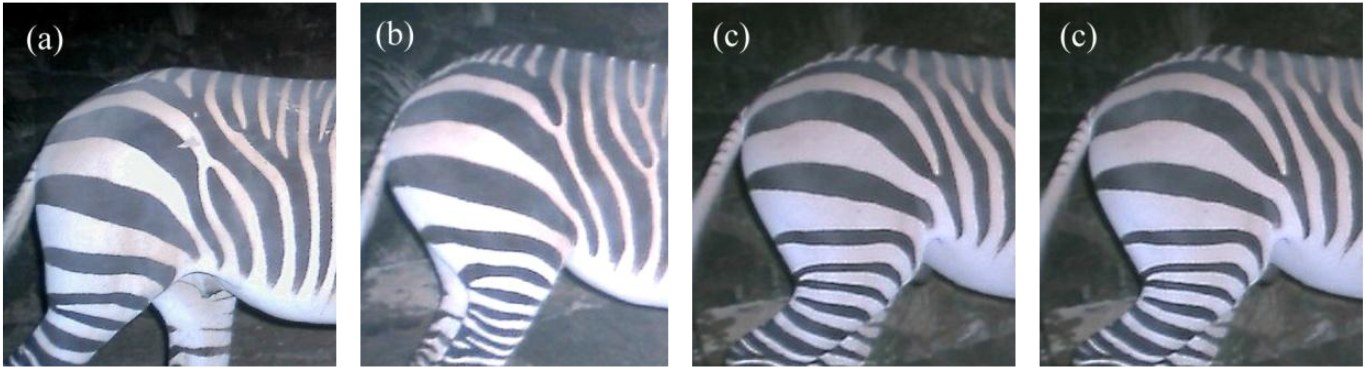
Variation in the gridiron stripe (for its location see Fig. 1). This stripe, from left to right, is (a) ‘simple’ – code G13, (b), ‘short’ - code G04, (c), joins 1^st^ rump stripe – code G02, (d) and ‘branched’ – code G05. The first body stripe (see Fig.1) is a ‘tree’ (a), ‘simple’ (b), ‘simple’ (c) and a ‘tree’ (d). All rump stripes in these examples are ‘simple’ (S13; see Appendix 1). Photos © L. M. Gosling.

The only rump stripes used are the two large stripes beneath the gridiron patch (Fig.1). The 1^st^ rump stripe is inserted just under the base of the tail. These stripes are unusual in being generally recognizable in fixed positions rather than relative to their neighbours. In this they resemble the gridiron and shoulder stripes and are coded in a slightly different way to the other stripes. In particular, where the 1^st^ body stripe joins the 2^nd^ rump stripe at their distal ends, forming a ‘V’ shape, I code these as S01 (i.e. 1^st^ rump stripe joining 2^nd^ rump stripe) rather than S07 (a ‘v’ shape in the case of all other body and rib stripes). However, with this exception, all stripes are coded in the same way for rump, body and rib stripes (see below) and a complete list of variants is given in Appendix 1.

Islands are black patches, of varying size and detached from the usual points of origin of stripes (the dorsal line of the body or the posterior edge of the rump). They are best illustrated by examples and one appears in Fig.2 (a) underneath the gridiron stripe. Islands are coded using their position relative to particular stripes and this system is restricted to the reference points of the gridiron stripe and the two adjacent rump and body stripes. The codes and definitions are given in Appendix 1.

All six main types of shoulder stripe are illustrated in Fig.3 (below) and listed in Appendix 1.

**Figure 3.**
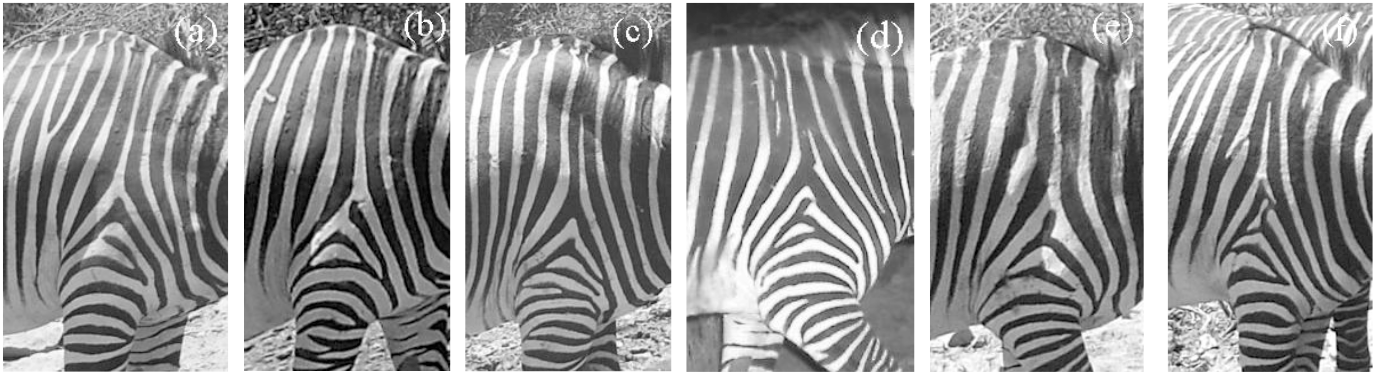
Examples of the six main types of shoulder stripe (see Fig.1). (a) and (b) are white stripes, respectively without (Sh1) and with (Sh2) a ‘triangle’ at the base. (c) and (d) are black stripes, respectively without (Sh3) and with (Sh4) a ‘triangle’ at the base. The shoulder stripe in Fig.1 is another example of Sh3. (e) and (f) are cases where the two black lines that border a white stripe join together before reaching the dorsal line; the two types are respectively without (Sh5) and with (Sh6) a ‘triangle’. In fact a ‘triangle’ includes any enclosed shape at the base of the stripe and in the example shown in (f) the shape is an irregular polygon. Photos © L. M. Gosling.

Apart from the gridiron and shoulder stripes, all stripes are coded using a unified approach. Examples are given in the legend to Fig. 1 above and codes and definitions are listed in Appendix 1.

The 13 characters (i.e. stripes plus islands) are established as columns in an Excel database with each row an individual zebra (Table 1).

The filter function in Excel (Data>Filter) is then used to search the database using the stripe variants defined from an animal in a new image. Depending on factors such as image quality, quality of the coded information in the ID library, distinctiveness of the individual in the new image, and so on, the number of candidate individuals (i.e. those not excluded by the filter process) are usually reduced to one or a few. When a single candidate individual is obtained it should always be checked against its reference photograph in the ID library. The time taken for the basic process of identifying a single candidate individual and checking against the candidate individual’s image an ID library of 3,156 individuals was 56+/-15 seconds (mean+/-SD) for 30 randomly selected, good quality, camera trap images. When candidates are filtered down to a few individuals these are each compared with the individual in the new image, initially using one distinctive character, then, when a likely individual was identified, with a number of additional characters from more than one part of the body. There is huge variation is all aspects of mountain zebra stripe patterns and when good images are available it is it is always possible to say with certainty whether or not an animal is the same as the one in the ID library.

Many animals and/or individual stripes unambiguously meet the standard definitions but others raise problems of definition. For example, when the character is one stripe joining another, the connection between two stripes may be indistinct. The problem in such cases is partly due to variable image quality: in a poor image a faint connection may appear to be absent so that a stripe appears to be a separate structure. There is also a point at which a ‘fork’ becomes a ‘branch’ and a problem of defining when a stripe is significantly shorter than its neighbours. In such cases I identified stripes as accurately as possible using the objective stripe criteria but was aware that a false negative identification was a possibility. The use of the filter function in Excel gives the opportunity to use stripes selectively. Thus if particularly conspicuous or unambiguous stripes occur on an image these can be used before ones that lack these characteristics; often animals can be identified before resorting to more ambiguous stripes. Also, if it is difficult to decide if a stripe is of one of two alternatives types, then both alternatives can be selected; this reduces the speed of identification but many possible animals will have been eliminated and a potential error avoided.

As the observer becomes familiar with the frequency of the variants in a population it is possible to select rare variants first which speeds up the identification process and reduces the number of stripes needed (which in turn reduces the potential for error). I explored the effect of this process by calculating the frequency of variants for all 13 characters in a randomly selected sample of 30 (good quality) images then compared the number of characters needed to identify each individual if the characters were filtered using either the order of the characters in the spreadsheet (see column headings above), a random sequence, and a sequence ranked by the frequency of each stripe variant at the particular position, in the population. The results in Table 2 show that selection in relation to frequency in the population can potentially speed up the process of identification by about 37% in relation to the list order and 44% in relation to the random order. While the size of these effects will be reduced by imperfect judgement about population frequencies of characters, the selection of rare variants by an experienced observer still makes an important contribution to the speed of identification.

**Table 2.**
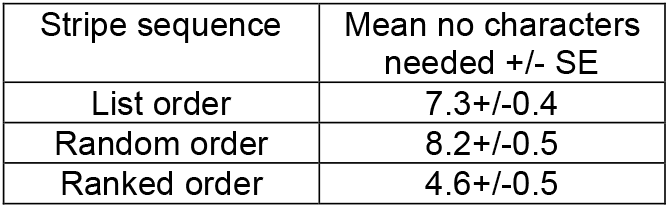
Comparison of the number of characters needed to identify individual mountain zebra when used in different sequences. The mean number needed when ranked according to the frequency in the population of a variant at a particular position is significantly less than both a random sequence and the order in the list given in the Excel table above (t = 5.62 and 4.40 respectively; p<0.001 in both cases).

When the search for an identity using the Excel filter is unsuccessful this may be either because the system has failed (false negative) or because the animal in the image is new. Failure may be because of some type of error (e.g. a coding mistake) or because the new image is poor or incomplete and does not allow accurate stripe coding. The next step is to repeat the search using different sets of characters (and different possibilities for ambiguous characters). I generally use two or three searches before concluding that an animal is probably new. The chance of this decision being correct is enhanced if only high quality images are used to set up the ID library and this is particularly important when studying large populations. Initially I adopted the procedure of searching through the entire reference library at this point, directly comparing each image with the new image. However, this process is becomes impractical when the ID library consists of more than, say, 2-300 individuals. For smaller numbers it is possible to do such checks using a localised stripe variant in the new image (one that is as rare and distinctive as possible) and then checking for it using a presence or absence approach as the reference library is scanned next to the new image. However with larger populations this is not practical and after two or three independent searches it must be assumed that the new image is of a previously unknown animal.

Whatever procedure is used it is certain that errors will be made and a few individuals will be entered into the library more than once. The search for such duplicates is a continuous process and in any ID search I generally checked all individuals selected when the number of alternatives are reduced to a small number. Duplicates are sometimes found in this way and can then be eliminated from the system. The main factor predicting the occurrence of duplicates are poor initial images which lead to errors in character coding. I recommend using only high quality images to set up IDs, that is, ones where all 13 characters can be coded with a high degree of certainty. I have adopted this criterion in recent study populations and in the most recent, the Naukluft NP study, the number of duplicates detected to date is 118 out of 5,335 individuals, or 1.5%; but of course this does not include any that are still undetected.

In some cases, individuals are seen so often that they become familiar to the observer and can be recognised on sight. Such recognition uses either a gestalt process or a few conspicuous characteristics or both. It may rely on the observer’s long term memory in some cases or on short term memory, for example when the same animal is drinking at a water hole in successive camera trap images. However, whenever an animal is recognised in this way it should always be checked against its image/s in the ID library. Memory plays tricks this final check is critical.

Simultaneous photography of right and left sides of the body is technically possible but for demographic studies the left and right sides of the body can be treated as separate populations. These ‘populations’ provide a check on some sampling biases. While they are clearly not truly independent, any disparity in, say, population estimates from right and left side populations indicates sampling problems such as differences in approach and departure routes past camera traps that need to be addressed. In practice, I used both sides in the first years Gondwana Canyon Park then switched to right sides for most routine analyses to save time when the population became large. I continued to collect some information about matching left sides when they provided important information about group membership for studies of social behaviour.

### POPULATION APPLICATION

The identification system described here can be used to monitor small or large populations of mountain zebra. As new animals are identified they can be added to a database list of individuals for a particular study site with columns for variables including year of birth (estimated for animals under two years old using developmental criteria described by Penzhorn, 1982b), gender, apparent relationships (mother and father, etc) and others depending on the objectives of the study. As the years pass, the proportion of the population that is of known age increases and, in the case of Gondwana Canyon Park, the known-age proportion of the population in the northern part of the Park that has been monitored since 2005, was over 70% by 2015. Such data allow demographic analysis including that of birth rates and age-specific survival and other information needed for age-structured modelling. The size of populations that can be routinely monitored are illustrated by the number of individuals identified. In 2025, the 6 main study sites where mountain zebra are routinely monitored contained over 12,000 coded individuals in the various ID libraries with about 7,800 known to be alive. The largest population, in the Naukluft extension of the Namib-Naukluft NP and adjacent properties contained over 5,200 coded individuals with over 3,200 animals known to be alive in 2018.

Where mountain zebra are relatively approachable and there is a good network of roads, it is also possible to monitor populations using normal photography. This is particularly useful in areas with large numbers of elephants because it is difficult to protect camera traps against these animals. One such area is Etosha NP where, in 2025, the ID library was up to 1,525 individuals with a maximum annual value of 940 known alive from retrospective census. I also used the individual recognition system to carry out a mark-recapture estimate of the mountain zebra population in this Park. Data were collected over seven days in August - September 2012 and 283 individuals were identified, mainly at six water holes (Jakkalswater, Renostervlei, Otjovasandu, Rateldraf, Klippan and Dolomietpunt). Using conventional mark-recapture formulae (Seber, 1982) and dividing the period into ‘mark’ and ‘recapture’ phases , these data yielded an estimate of 802+/-116 mountain zebra (Gosling, 2012). This compared with an estimate of 685+/-158 from an MET air survey carried out earlier in August (Kolberg, 2012).

## DISCUSSION

Key issues in individual recognition are the reliability of each identification and the speed of the process. To ensure reliable identification any new image should always be checked by direct visual comparison with reference photograph/s of the candidate individual and that confirmation should involve at least 3 variable characters. This requirement places a time constraint on the identification process but is unavoidable to avoid errors including the creation of duplicates in the reference library which are time consuming to remove. This time constraint applies to both manually coded and computerised pattern recognition and is absent only in small populations where direct visual comparison is the primary recognition technique.

The main alternative to the manual coding system described here are computer-based systems of pattern recognition using an approach related to the well-established process of human fingerprint recognition and undergoing rapid advance with current developments in machine learning. Progress in this field includes the pattern recognition of plains zebra (Foster et al., 2006) and increasingly powerful computerised photographic identification systems for plains and Grevy’s zebra (‘StripeSpotter’, Lahiri et al., 2011; ‘Hotspotter’, Crall et al., 2013 and Stennett et al., 2022). Both systems probably depend to some extent on the experience of the user although computerised pattern recognition system may be preferable where only a small investment of time for training is possible at the start of the project and where project staff work for relatively short periods. In terms of the time involved in identification for an individual zebra, once an observer is trained, the systems are probably similar. Time benefits must accrue when pattern recognition systems become fully automatic (e.g. Sherley et al., 2010 and Stennett et al., 2022) but perhaps at the cost of losing contact with the natural history of the subjects and an unknown risk of false negatives. The advantages of the manual coding system include the fact that incomplete and oblique images can often be used (although Stennett et al., 2022, have started to address the problem of including 3-D images in computerised identification) and that because of the level of scrutiny needed the observer benefits from a close familiarity with the study animals and the opportunity to collect additional information such as that on condition and scars from fighting and failed predation attempts.

Whatever the system used, the issue of false negatives should be dealt with and the manual coding system described here is designed partly with this in mind. If a new image is not identified by using the filtering system, then alternative sequences of characters can be used and this property of the system reduces the chance of false negatives. The option of visually checking by searching the entire reference library become impractical when populations are large. However, since the assumption that any animal is new can only be probabilistic, adding an animal as new without checking all existing images will inevitably result in some animals being given two reference identities. Strategies for avoiding such false negatives IDs include restricting the ID library to high quality photographs in which all 13 characters can be coded unambiguously and adopting conservative filtering approaches, such as using more than one code at a stripe position if there is any doubt about the type of stripe. It is also best to routinely check the reference photographs of small groups of individuals when the filtering process reveals animals with similar stripe codes, rather than always trying to reduce the alternatives to one individual.

Camera trapping has developed rapidly over the past decade and is an excellent technique for individual-based studies. Mountain zebras are water-dependent and this provides an opportunity for placing camera traps in locations that potentially sample entire populations. An additional benefit of camera trapping is that the accumulated photographs (which should be carefully archived) provide a rich resource for further study. Photograph libraries can be re-sampled for different practical and academic purposes. For example samples can be sampled to code for individual-based variation in body condition to compare responses to variation in the food supply and condition-dependent changes in behaviour; and identities and patterns of association can be used for the investigation of social group membership in studies of life-history evolution.

## ACKNOWLEDGEMENTS

I am grateful to the owners and staff of Gondwana Canyon Park, the NamibRand Nature Reserve, Büllsport Guest Farm, the Neuras Wine and Wildlife Estate, Ababis Guest Farm, the Namib Naukluft Lodge and Schlesien Farm for access and support while the technique reported here was developed and employed and for helping look after and, in some cases, supplying, camera traps. Chris Brown originally suggested the study of mountain zebra and agreed to host the Mountain Zebra Project at the Namibia Nature Foundation. The data used from Gondwana Canyon Park for recent analyses including the ‘time taken’ analysis were collected by Michelle Rodgers and her field team. Thanks also to the Ministry of Environment, Forestry and Tourism, especially Kenneth Uiseb, Manie le Roux, Werner Killian and Shayne Kötting for support and access to National Parks including the Ai-Ais National Park, the Namib-Naukluft National Park, Etosha National Park and the Hobatere Concession Area; and the National Commission for Research, Science and Technology for permission to carry out research in Namibia most recently under Authorization No. AN20180715. The Rufford Foundation, the Whitley Fund for Nature, the GaiaZOO and the Parc Zoologique de Montpellier provided generous financial support and the Namibia Nature Foundation and Newcastle University provided administrative support.

## Appendix A

Stripe categories used for individual recognition.

Gridiron stripe variants (see Figs 1 and 2):

| Code | Description |
| --- | --- |
| G01 | The gridiron stripe joins the 1<sup>st</sup> body stripe. |
| G02 | The gridiron stripe joins the 1<sup>st</sup> rump stripe. E.g. Fig.2 (c). |
| G03 | The 'stripe' part of the character (i.e. the linear section that runs obliquely down from the 'gridiron' section at the top) is entirely missing. |
| G04 | The 'stripe' part of the character is reduced to length that is distinctly shorter than normal and (usually) than the body stripes. It may be up to $\frac{3}{4}$ the normal length or a very small stub. E.g. Fig.2 (b). |
| G05 | A large branch e.g. Fig.2 (d); nearly always 2 branches but if more they are included here. |
| G06 | Does not exist – listed here for consistency with other stripes. |
| G07 | Does not exist – listed here for consistency with other stripes. |
| G08 | Does not exist – listed here for consistency with other stripes. |
| G09 | A short 'spur' projects from the side of an otherwise simple stripe. |
| G10 | A white 'needle eye' in an otherwise simple stripe. |
| G11 | A small fork at the end of an otherwise simple stripe. |
| G12 | Does not exist – listed here for consistency with other stripes. |
| G13 | A simple linear structure with a pointed or rounded end. E.g. Fig.1 and Fig.2 (a). |

Islands: codes and definitions:

| Code |  |
| --- | --- |
| Is1 | No islands in the positions defined for Is2-6. |
| Is2 | Island beneath gridiron stripe (and above 1<sup>st</sup> rump stripe). E.g. Fig.2 (a). |
| Is3 | Island beneath 1<sup>st</sup> rump stripe. |
| Is4 | Island beneath 2<sup>nd</sup> rump stripe. |
| Is5 | Island in front of gridiron stripe |
| Is6 | Island in front of 1<sup>st</sup> body stripe. |

Shoulder stripe variants and their codes:

| Code | Description |
| --- | --- |
| Sh1 | The stripe is a white vertical bar approximately aligned with the centre of the foreleg. It is between two vertical black lines that converge from in front of and behind the top of the foreleg. E.g. in Fig.3 (a). No triangle (see below). |
| Sh2 | The stripe is similar to Sh1 but includes a 'triangle' at the base. E.g. in Fig.3 (b). Other examples of triangles in other shoulder stripe types can be seen in Fig.3 (d) and (f). |
| Sh3 | The stripe is a vertical black line approximately aligned with the centre of the foreleg. No triangle. E.g. Fig.3 (c) and Fig.1. |
| Sh4 | The stripe is similar to Sh3 but includes a 'triangle' at the base. E.g. in Fig.3 (d). |
| Sh5 | The stripe resembles Sh1 but the two vertical bordering lines join up before reaching the top of the body. E.g. Fig.3 (e). |
| Sh6 | The stripe is similar to Sh5 but includes a 'triangle' at the base. E.g. in Fig.3 (f). As mentioned in the figure legend 'triangles' include almost any enclosed shape and in the case shown in Fig.3 (f) the shape is an irregular polygon. |

Stripes: codes and definitions. These apply to all stripes except gridiron and shoulder stripes:

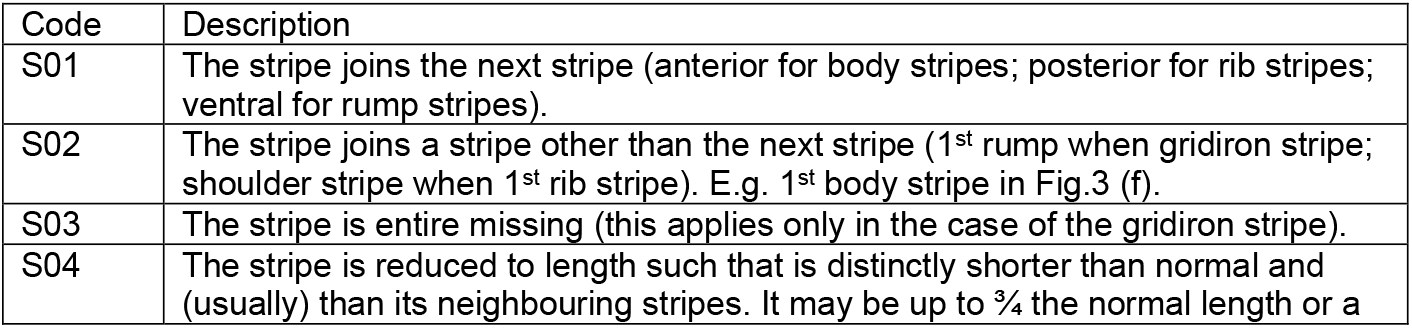
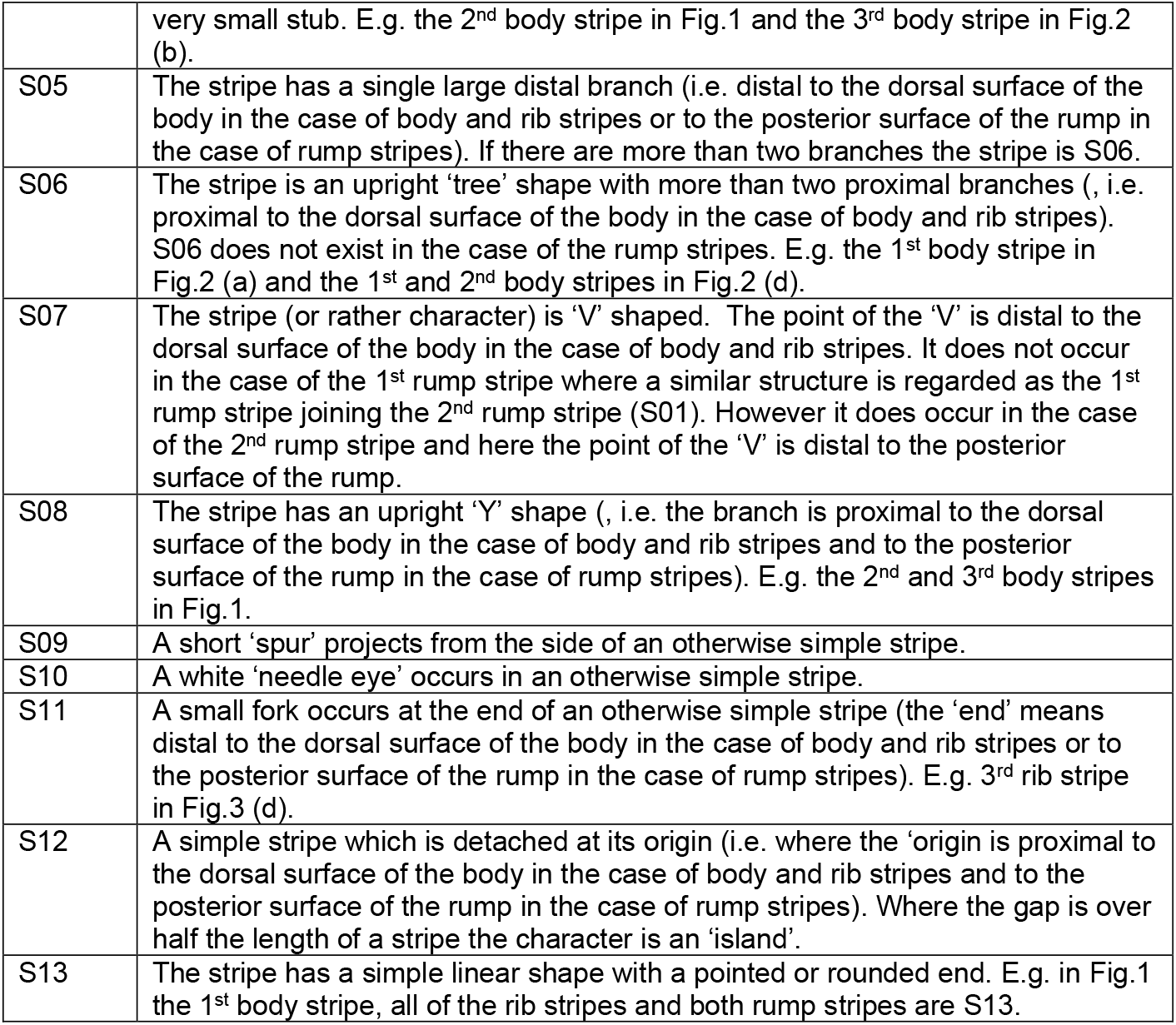

